# Spatial dynamics of IFITM1: a core component of the interferon-stimulated gene-resistance signature in glioma

**DOI:** 10.1101/2025.07.16.661931

**Authors:** Yolanda Solans Bara, Erisa Nita, Zuzana Gabrhelikova, Olimpia E Curran, Paul M Brennan, Ashita Singh, Kathryn L Ball

## Abstract

The IFITM1 protein is a key component of the Interferon-Stimulated Gene (ISG) network, which has been linked to treatment-resistant signatures in various cancers, including glioblastoma (GBM). Despite its impact, the mechanisms underlying IFITM1’s role in cancer remain poorly understood. Here, we demonstrate that the spatial dynamics of IFITM1 localisation are highly context-dependent, particularly in GBM tissue. In the vasculature, IFITM1 is predominantly localised to the plasma membrane of endothelial cells. However, when present in a subset of cancer stem cells, IFITM1 adopts a distinct perinuclear location, where it co-localises with IFITM3 and is notably absent from the membrane. The spatial dynamics of IFITM1 localisation was investigated in patient-derived glioma stem cells (GSCs), primary endothelial cells and engineered cell lines. Endogenous IFITM1 localised primarily to assemblies in the perinuclear space, however loss of functional IFITM3 led to a shift in the distribution of the protein to a predominantly membrane location. In contrast, IFITM3 localisation was unaffected by the loss of IFITM1. The results were recapitulated by transient expression of IFITM1 or -3 into double knockout (DKO) GSCs and by using an engineered system where IFITM1 expression was IFN-independent. Co-expression studies demonstrate that IFITM3 is sufficient to localise IFITM1 to vesicle structures in the perinuclear space and that mutant forms of IFITM3 lead to retention of IFITM1 primarily at the plasma membrane. Our data highlights dynamic changes in the subcellular localisation of IFITM1 suggesting that distinct functions of this resistance factor may present a specific target for therapeutic intervention.

## Introduction

Glioblastoma (GBM) presents a significant clinical challenge as the most aggressive and invasive form of brain cancer, with a median life expectancy of approximately 15 months following diagnosis. Despite advances in treatment, current therapeutic approaches—including surgical resection, radiation therapy, and chemotherapy—have limited success in improving patient outcomes. Due to the highly infiltrative nature of GBM, complete surgical removal is unachievable, leading to frequent recurrence, most commonly at the primary tumour site. The prevailing hypothesis suggests that GBM relapse is driven by a population of treatment-resistant cancer stem cells that persist after treatment, and that exhibit strong resistance to standard of care therapies (Eckerdt and Platanias 2023; Shi et al. 2023; Tang et al. 2021).

Other than treatment-resistant stem cell populations, angiogenesis plays a critical role in GBM therapeutic resistance, as well as its pathophysiology (Ahir, Engelhard, and Lakka 2020; Rahman and Ali 2024). GBM are characterised by a highly vascularised microenvironment, driven by hypoxia-induced secretion of pro-angiogenic factors, including vascular endothelial growth factor (VEGF) (Mosteiro et al. 2022; Liu et al. 2023). These factors activate signalling pathways that promote endothelial cell proliferation, migration, and tube formation. Additionally, aberrant expression of other angiogenic mediators contributes to vascular remodelling and stabilisation. This extensive angiogenic activity supports the metabolic demands of the rapidly growing tumour, facilitating its invasiveness and resistance to conventional therapies.

Interferon-induced transmembrane protein 1 (IFITM1) was first identified for its role in the host’s antiviral defence but has since gained attention for its involvement in processes such as proliferation, invasion, and apoptosis (Friedlova et al. 2022; Gomez-Herranz, Taylor, and Sloan 2023). As a well-characterised Interferon-Stimulated Gene (ISG), IFITM1 has also emerged as a key factor in resistance to multiple cancer treatments, forming part of the ISG-Resistance Signature (ISG-RS) (Weichselbaum et al. 2008; Benci et al. 2019). Additionally, it plays a role in maintaining “stemness” and contributes to stem cell resistance to interferons (IFNs) and viral infections (Friedlova et al. 2022; Wu et al. 2018). Both IFITM1 and its close relative, IFITM3, have been implicated in angiogenesis across various cancers, including glioma (Xiong et al. 2024; Popson et al. 2014). Multiple lines of evidence—such as gene expression studies, functional assays, and in vivo models—suggest that these proteins promote angiogenesis by enhancing endothelial cell functions (Popson et al. 2014; Xiong et al. 2024; Friedlova et al. 2022). As a result, their modulation influences blood vessel formation, particularly in pathological conditions like tumour growth. However, the precise molecular mechanisms by which IFITM1 and IFITM3 contribute to angiogenesis remain unclear.

IFITM1 has been implicated as a potential driver in a diverse range of cancers, including GBM. Transcriptomic analyses indicate that the ISG-RS is upregulated in tumours from at least 40% of patients with high-grade gliomas, which includes GBM (Weichselbaum et al. 2008). Thus, IFITM1 holds promise as both a biomarker and a potential therapeutic target. Despite these insights, the specific mechanisms through which IFITM1 contributes to cancer pathogenesis, particularly in GBM, remain to be fully elucidated. Here we show that IFITM1 exhibits distinct subcellular localisation patterns in different cell types within GBM tissue. While IFITM1 is detected in the membranes and cytoplasm of epithelial cells within the tumour vasculature, its distribution within glioma tumour cells differs significantly. Specifically, IFITM1 is localised to the perinuclear space in a subpopulation of tumour cells, where it is absent from the plasma membrane. To investigate the mechanisms governing IFITM1 subcellular dynamics, we utilised patient-derived glioma stem cells (GSCs) and an orthogonal cervical cancer cell model. We confirmed that endogenous IFITM3 and IFITM1 interact and demonstrate that the loss of IFITM3 function resulted in a redistribution of IFITM1, undercovering a role for IFITM3 in determining IFITM1 localisation. Moreover, a mutant form of IFITM3 demonstrated that it is sufficient to direct IFITM1 to specific subcellular location, whereas IFITM1 status did not influence the localisation of IFITM3. Given the critical role of cancer stem cells in glioma self-renewal and rapid tumour growth, as well as the highly vascularised nature of GBM, our findings highlight the potential of IFITM1 subcellular localisation as a specific target for therapeutic intervention in GBM.

## Results

### IFITM1 and IFITM3 have distinct subcellular localisations in cells within glioma tissue

Determining a protein’s spatial location in its native cellular or tissue context, provides detailed insights into how it is organised and regulated with respect to other factors and whether the local environment impacts on its function. Notably, whereas IFITM3 is reported to be in late endosomes, IFITM1’s antiviral activity is primarily linked to its presence at the plasma membrane, where it is thought to alter lipid bilayer composition and fluidity, contributing to the prevention of virus particle entry (Guo et al. 2021; Chesarino et al. 2017; Weston et al. 2014). Despite extensive evidence that IFITM1 and IFITM3 contribute to drug resistance and angiogenesis in some cancers, their mechanism(s) of action in this diseases state are not fully understood (Friedlova et al. 2022; Rajapaksa, Jin, and Dong 2020). To address this gap in our knowledge, we initiated a study on the spatial expression of the IFITM1 protein relative to IFITM3 in cancer tissue.

Since both IFITM1 and IFITM3 have been implicated as potential drivers in glioma (Yu et al. 2011; Xiong et al. 2024), we analysed their endogenous distribution and subcellular localisation in GBM tissue. Immunohistochemistry (IHC) was performed on cores from a proprietary tissue microarray (TMA) generated using tumour tissue from patients with a range of brain cancers (Al Shboul et al. 2024). IFITM1 showed strong staining around the vasculature in primary GBM (Grade IV) (Fig. 1A). To identify the specific cell types expressing IFITM1, we employed tissue immunofluorescence (IF) and co-staining with cell-type-specific markers. The IFITM1-positive, vasculature-associated cells were also positive for the endothelial cell marker CD31 (Fig. 1B), and negative for the stem cell marker SOX2 (Fig. 1C). Thus, IFITM1 is present in CD31-positive endothelial cells, where it is closely associated with CD31 at the plasma membrane, although only limited colocalisation was observed (Fig. 1C; yellow signal).

**Figure 1.**
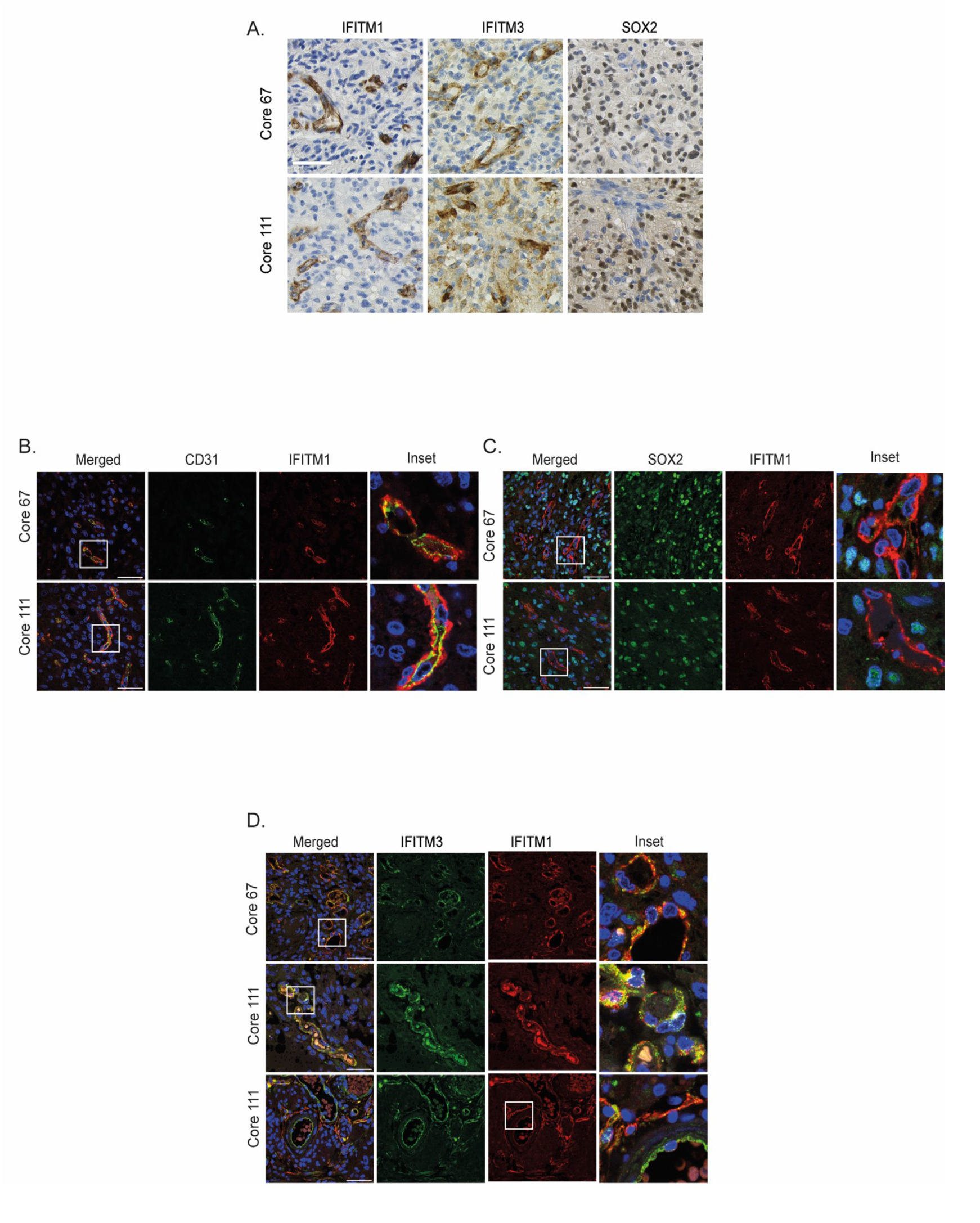
IFITM1 and IFITM3 colocalise at the plasma membrane of TECs in glioma tissue. (**A**) IHC of tissue cores from 2 patients with Grade 4 GBM (IDH1-WT). The sections were stained individually with anti-IFITM1, -IFITM3 or -SOX2 antibodies and developed using DAB-staining and counterstained with haematoxylin. Scale bar = 50 µm (**B-D**) Confocal images of two cores (as above) were stained using anti-IFITM1, -IFITM3, -CD31 or -SOX2 as indicated, plus DAPI. Scale bar = 50 µm; n = 3.

Consistent with recent reports linking IFITM3 to angiogenesis in GBM, IHC using an IFITM3-specific antibody also revealed strong staining around blood vessels (Fig. 1A). Perhaps more surprisingly, given that IFITM3 is often reported as a predominantly endosomal protein (Guo et al. 2021; Chesarino et al. 2017), immunofluorescence analysis demonstrated a high degree of co-localisation between IFITM1 and IFITM3 at the plasma membrane of endothelial cells (Fig. 1D; yellow signal, top two panels). However, there were also regions within the vasculature where membrane staining of IFITM1 and IFITM3 remained distinct, indicating potential spatial differences in their membrane distribution (Fig. 1D; bottom panel).

In contrast to the endothelial cells of the vasculature where IFITM3 had a plasma membrane localisation, IFITM3 showed a distinctive pattern of condensed perinuclear staining in glioma tissue tumour cells (Fig 2A) by IHC. Immunofluorescence confirmed the distribution of IFITM3 within the perinuclear region of a population of individual tumour cells (Fig 2B). Further, IFITM3 positive tumour cells were generally seen to be positive for SOX2 (Fig 2B) suggesting that IFITM3 resides within the stem cell niche. Using IF, we were also able to detect IFITM1 in a subset of the IFITM3 positive stem cells. When IFITM1 was present in the stem cell population it was colocalised with IFITM3 in condensed perinuclear assemblies and was noticeably absent from the plasma membrane (Fig. 2C). Thus, IFITM1 has a dynamic spatial distribution within the GBM tissue cells with predominantly membrane staining in the endothelial cells and perinuclear staining in a subpopulation of IFITM3 stem cells.

**Figure 2.**
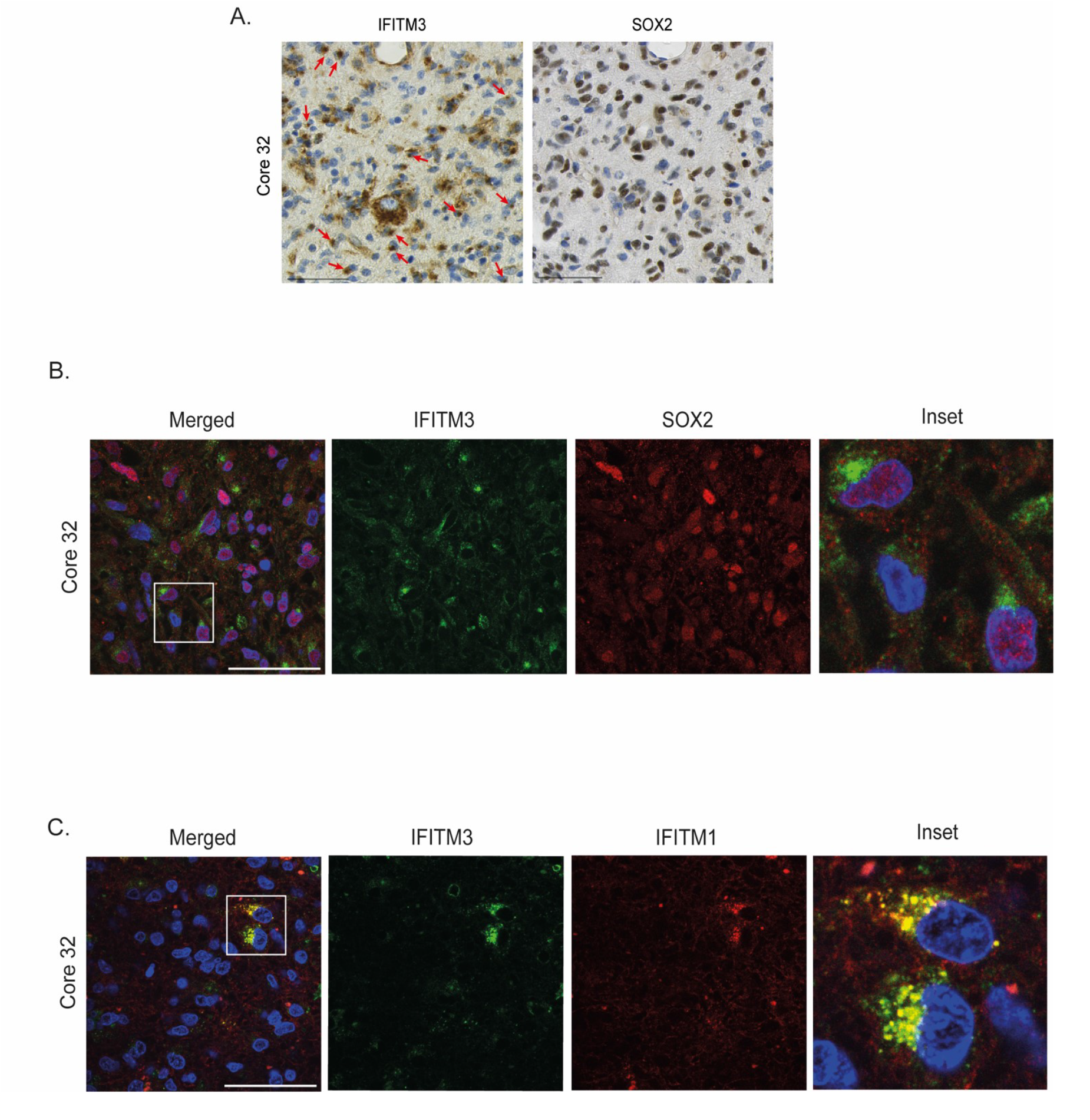
IFITM1 and IFITM3 colocalise in the perinuclear space of SOX2 positive cancer cells. (**A**) IHC of a tissue core from a patient with Grade 4 GBM (IDH1-WT). The sections were stained individually with anti-IFITM3 or -SOX2 and developed using DAB-staining and counterstained with haematoxylin. Scale bar = 50 µm (**B, C**) Confocal images of a glioma core stained with anti-IFITM3 and -SOX2 antibodies (**B**) or anti-IFITM1 and -IFITM3 antibodies (**C**) as indicated, plus DAPI. Scale bar = 50 µm; n = 3.

### IFITM1 and IFITM3 are perinuclear in GSCs and primary endothelial cells

Most studies indicate that IFITM1 is primarily localised at the plasma membrane as seen in glioma endothelial cells, we were therefore curious about its perinuclear localisation in some GBM tissue tumour cells. To investigate the distribution of IFITM1 in greater detail, we utilised cell systems. Two patient-derived GSC lines (E20 and E27), both of which exhibited detectable levels of endogenous IFITM1 and IFITM3 protein in response to IFNA2 treatment (Fig. 3A), were analysed by fluorescence microscopy. Figure 3 demonstrates that IFITM1 (red) and IFITM3 (green) were essentially co-located (as indicated by the yellow signal in the merged image; Fig 3B and C) in both GSC lines. Furthermore, consistent with the spatial distribution observed in GSCs within GBM tissue (Fig. 2), both IFITMs were primarily localised to the perinuclear space, with little to no plasma membrane staining detected.

**Figure 3.**
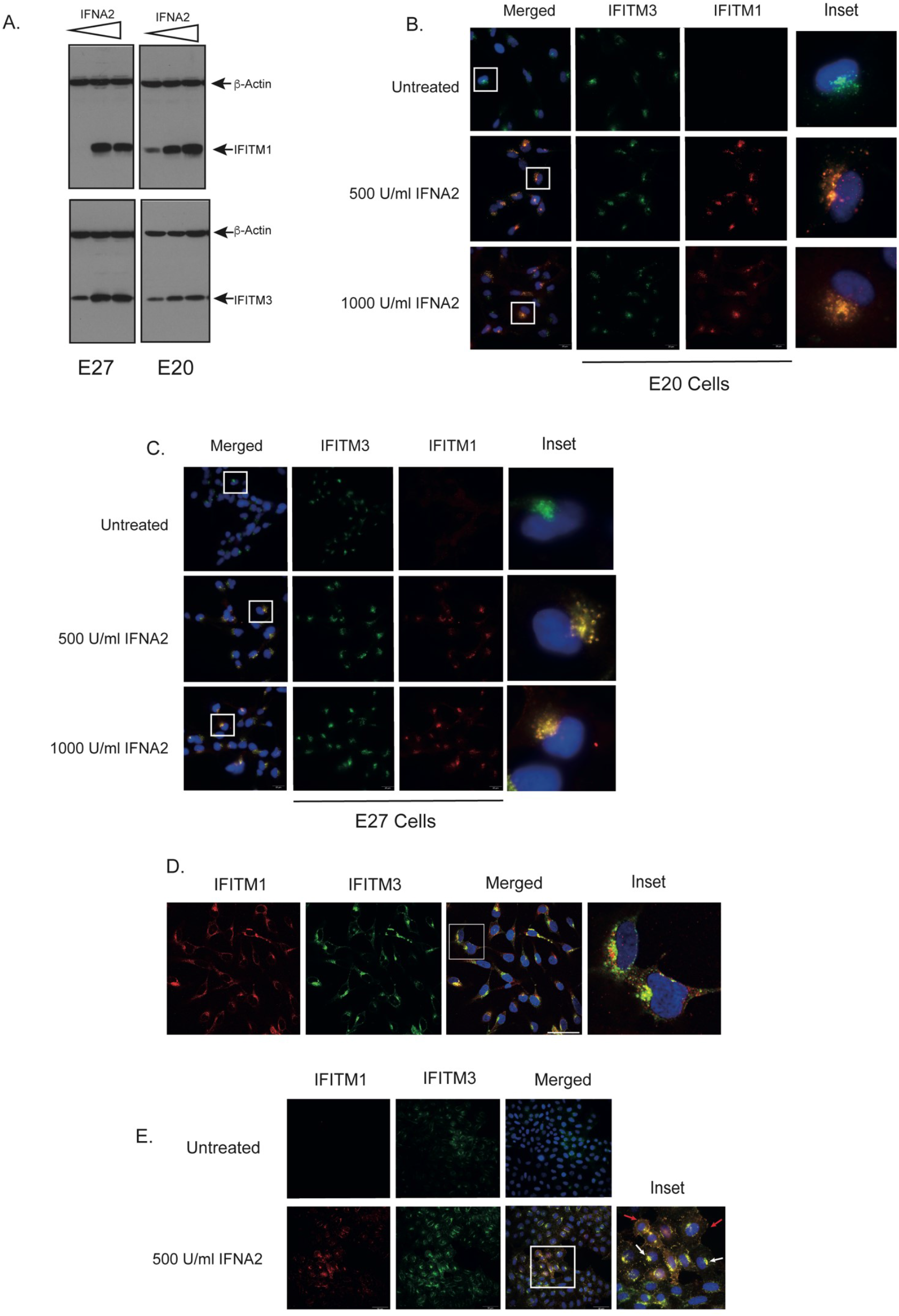
IFITM1 colocalises with IFITM3 in perinuclear assemblies following IFNA treatment. (**A**) Immunoblots of E20 and E27 GSC lysate from control and IFNA-treated cells (500 and 1000 U) 24 h post-treatments. The immunoblots were developed using anti-IFITM1 or anti-IFITM3 antibodies then overlayed with anti-β-actin antibody. Immunofluorescence of E20 (**B**) and E27 (**C**) GSCs stimulated for 24 h with IFNA (500 and 100 U) or the carrier alone. IFITM1 was detected using Alexa 549 (red) secondary antibody and IFITM3 was detected using Alexa 488 (green) secondary antibody. The nucleus was stained with DAPI (blue). Scale bar = 25 μm; n = 3. (**D**) Immunofluorescence of untreated HCMEC/D3 where IFITM1 and IFITM3 were detected as above. The nucleus was stained with DAPI (blue), and the scale bar is 100 μm. (**E**) Immunofluorescence of SiHa cells incubated for 24 h with IFNA (500 U) or the carrier alone. IFITM1 and IFITM3 were detected as above. The nucleus was stained with DAPI (blue). Scale bar = 50 μm; n = 3.

In contrast to GBM tissue, where both IFITM1 and IFITM3 were localised at the plasma membrane of endothelial cells lining the blood vessels, isolated primary human blood-brain barrier (BBB)-derived endothelial cells, like GSCs, exhibited a perinuclear localisation for both proteins (Fig 3D). This difference in subcellular localisation between endothelial cells in culture and those in the tissue vasculature suggests that the distribution of IFITM1 and IFITM3 is spatially dynamic and that it is determined not solely by cell lineage, but also by the local tumour environment and/or by the nature of tumour-associated endothelial cells compared to those not associated with the tumour.

We have previously used the cervical cancer cell line (SiHa) to study the role of IFITM1 in mRNA translation (Gomez-Herranz et al. 2019) and were interested to know where IFITM1 and IFITM3 were localised in this well-established tumour cell line compared with patient-derived GSCs and primary BBB endothelial cells. In unstimulated SiHa cells, where IFITM1 was barely detectable, IFITM3 was localised in perinuclear assemblies (Fig 3E) as seen in GSCs (Fig 3B & C). Following IFNA2 treatment, IFITM1 and IFITM3 were again primarily colocalised in perinuclear assemblies, however unlike the GSCs there was also some co-localisation detectable at the plasma membrane. Thus, endogenous IFITM1 can localise to the perinuclear space and in some cells to the plasma membrane highlighting the dynamic nature of the protein.

### IFN is not the main determinant of IFITM1 localisation

As IFN-treatment was required to detect IFITM1 in the GSC and SiHa backgrounds, we were not able to determine how IFITM1 localisation was influenced by IFNA2. In other words, would IFITM1 be primarily plasma membrane-associated in the absence of IFNA and is perinuclear localisation a result of IFN-activated post-translational regulation of IFITM1? To answer this question, we noted that primary endothelial cells had high basal levels of endogenous IFITM1, which colocalised with IFITM3 in perinuclear assemblies (Fig 3E) in the absence of IFN treatment. As with SiHa cells (Fig 3D), a portion of the IFITM1 in the BBB endothelial cells was seen at the plasma membrane although most of the protein was still seen in perinuclear assemblies. Next, we used an orthogonal system where IFITM1 was expressed using the Flp-In system integrated into SiHa cells in which IFITM1 had been knocked-out using the RNP gene-editing method.

The Flp-In:IFITM1 cells had basal levels of IFITM1 protein expression similar to that seen in IFNA2-stimulated wild-type (WT) SiHa cells (Fig 4A) providing an IFN-independent system to study IFITM1 localisation. Notably, the levels of IFITM1 decreased slightly after IFN treatment, suggesting that it increases exogenous IFITM1 turnover. When the localisation of IFITM1 in the Flp-In:IFITM1 cells was examined, though it was more dispersed than in IFNA2 treated WT cells (Compare Fig 4B: Flpin:IFITM1 untreated to Fig 3E WT treated), the majority of the staining was perinuclear and in most cells it was associated with IFITM3 (Fig 4B; Flp-in:IFITM1 untreated). When these cells were treated with IFNA2, IFITM1 became more concentrated in the perinuclear assemblies, though it should also be noted that some cells showed IFITM1 on the plasma membrane. Thus, exogenous IFITM1 can localise to the perinuclear space in the absence of IFNA but that IFNA2 signalling promotes a more condensed distribution. Notably, in IFNA2 treated cells, exogenous IFITM1 localisation mirrored that of endogenous IFITM3 (Fig 4).

**Figure 4.**
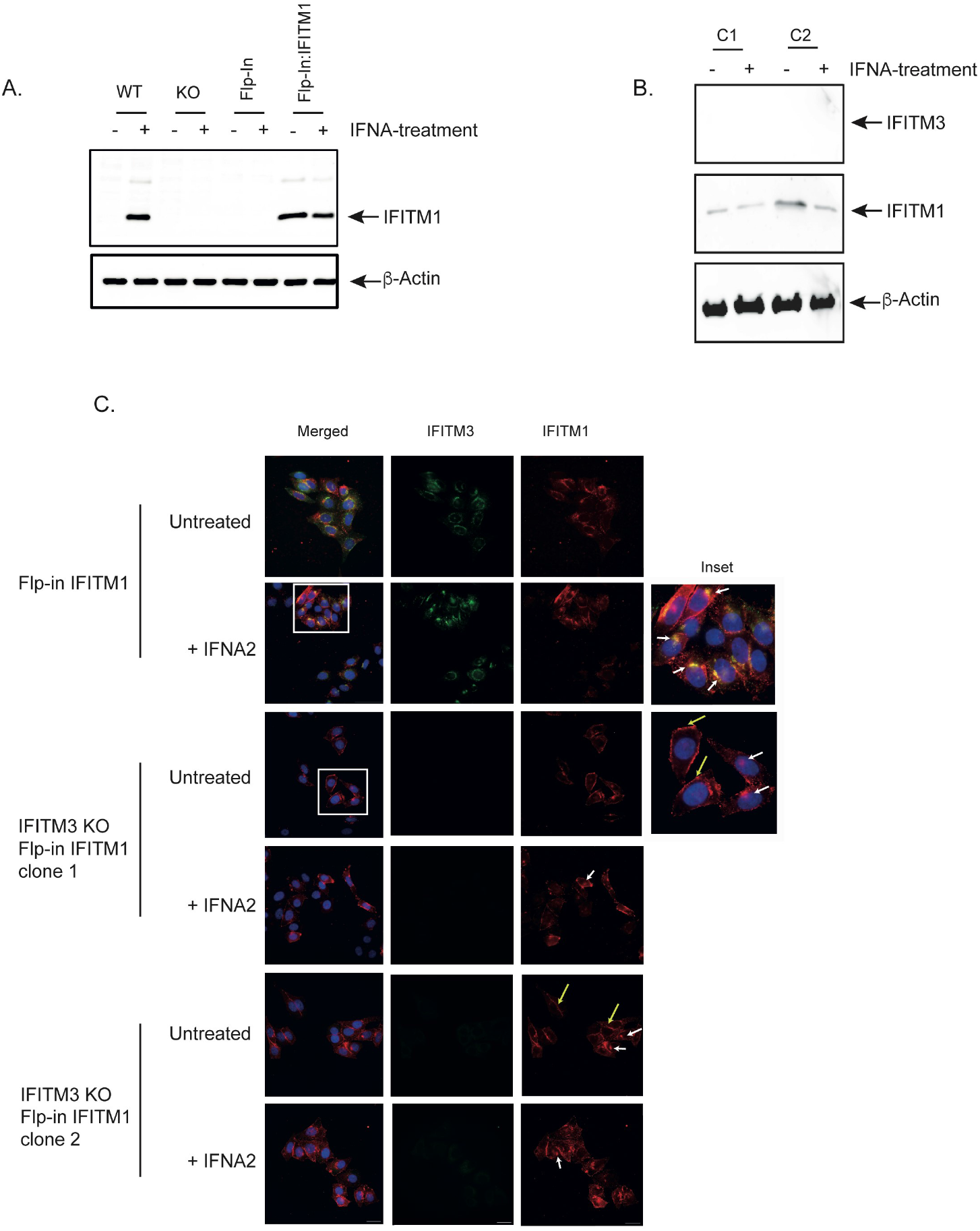
Loss of IFITM3 affects the localisation of exogenous IFITM1. (**A**) Immunoblot of lysates from WT, IFITM1 KO and control IFITM1 KO Flp-In SiHa cells (Flp-In) plus IFITM1 KO Flp-In stably expressing exogenous IFITM1 (Flp-in:IFITM1) untreated or treated for 24 h with IFNA (500 U). The immunoblot was developed using anti-IFITM1 antibody, a separate gel was used to analyse β-actin levels. (**B**) Flp-in:IFITM1 SiHa cells were used to generate IFITM3 KO immunoblot showing two independent clones (C1 and C2) treated with IFNA (500 U) or carrier-alone for 24 h prior to harvesting. The immunoblots were developed using anti-IFITM1 or anti-IFITM3 antibodies. The IFITM3 immunoblot was subsequently overlayed with anti-β-actin antibody. (**C**) Immunofluorescence of Flp-in:IFITM1 SiHa cells and 2 IFITM3 KO clones (1 and 2) in the Flpin:IFITM1 background. Cells were incubated for 24 h with IFNA (500 U) or the carrier alone. IFITM1 was detected using Alexa 549 (red) and IFITM3 was detected using Alexa 488 (green) secondary antibody. The nucleus was stained with DAPI (blue), and the scale bar is 50 μm.

Next, we generated two independent IFITM3 KO lines (Figure 4B and C; clone 1 and 2) in the Flpin:IFITM1 background. The first thing of note was that there was more IFITM1 on the plasma membrane in untreated IFITM3 KO Flp-in:IFITM1 SiHa cells than in the untreated Flp-in IFITM1 control (Fig 4B; yellow arrows). Additionally, a subset of cells showed weak perinuclear IFITM1 staining (Fig 4B; white arrows). Although there were some alterations in IFITM1 distribution following IFNA2 treatment (the appearance was more granular), there was no significant difference in the frequency of perinuclear staining between the treated and untreated cells.

Data from endothelial cells and the orthogonal Flp-In:IFITM1 cells suggests that activation of IFN signalling is not required for IFITM1 to localise to perinuclear assemblies. In addition, the status of IFITM3 appears to be a factor in determining the distribution of exogenous IFITM1 in the FlpIn:IFITM1 SiHa model, though we cannot entirely rule out a contribution from additional IFNA-dependent factors.

### The dynamics of IFITM1 localisation is regulated by IFITM3

To dig deeper into the potential role of IFITM3 in the regulation of IFITM1 localisation, we turned to an endogenous IFITM1 and IFITM3 system using GSCs. First, individual IFITM1 and IFITM3 KO lines were generated in the GSC E27 background so we could ascertain if the localisation of either of these factors influenced that of the other. Though the level of IFITM3 protein in the IFITM1^-/-^ GSCs treated with IFNA was reproducibly lower than in the WT background (Fig 5A), there was no obvious difference in its localisation (Fig 5B; compare IFITM3 (green) in IFITM1 KO and WT lines). Strikingly however, the loss of IFITM3 function (Fig 5B; IFITM3 KO) led to a marked redistribution of IFITM1. Thus, in the absence of IFITM3 (IFITM3 KO), rather than being concentrated in the perinuclear space (see WT panels), IFITM1 displayed predominantly membrane staining accompanied by a more diffuse punctate cytoplasmic distribution pattern.

**Figure 5.**
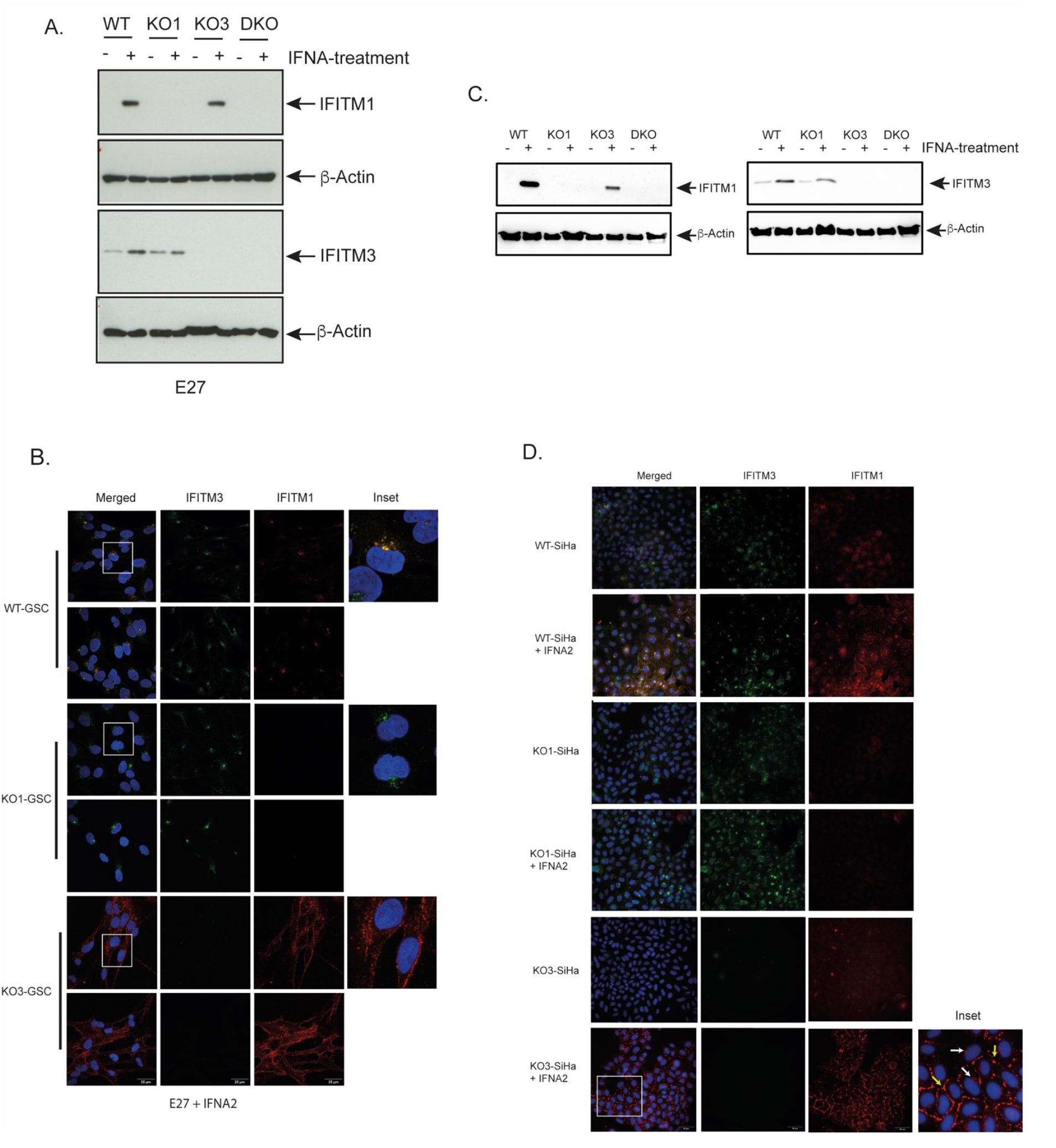
Loss of IFITM3 impacts the localisation of endogenous IFITM1 across different cell models. (**A**) Immunoblot analysis of IFITM1 KO (KO1), IFITM3 KO (KO3), and double KO (DKO) lines generated in the E27 GSC background following treatment with IFNA (500 U) or carrier alone for 24 h. Membranes were probed with anti-IFITM1 and anti-IFITM3 antibodies and IFITM1 membrane was re-probed with anti-β-actin as a loading control. (**B**) Immunofluorescence staining of WT E27 GSCs (WT-GSC), IFITM1 KO (KO1-GSC) and IFITM3 KO (KO3-GSC) after incubation with IFNA (500 U) for 24 h. IFITM1 was detected using Alexa 549 (red) and IFITM3 was detected using Alexa 488 (green) secondary antibody; nuclei were stained with DAPI (blue). Representative images from two fields per condition are shown (n = 3). Scale bar: 25 μm. (**C**) Immunoblots of WT, IFITM1 (KO1) and IFITM3 (KO3) KOs SiHa cells treated with IFNA (500 U) or carrier for 24 h. Membranes were probed with antibodies against IFITM1 or IFITM3, and the IFITM1 membrane was subsequently overlayed with anti-β-actin antibody. (**D**) Immunofluorescence of WT SiHa (WT-SiHa), IFITM1 KO (KO1-SiHa) and IFITM3 KO (KO3-SiHa) after incubation with IFNA (500 U) or carrier for 24 h. IFITM1 was detected using Alexa 549 (red) and anti-IFITM3 was detected using Alexa 488 (green) secondary antibody. The nucleus was stained with DAPI (blue). Scale bar = 50 μm; n = 3.

To independently validate the findings observed in GSCs, the SiHa cell model was employed and individual IFITM1 and IFITM3 KO lines were analysed. Again, although IFNA treatment led to a reduction in IFITM3 levels in IFITM1 KO SiHa cells (Fig. 5C), the data demonstrated that IFITM3 was still able to localise to perinuclear assemblies in the absence of IFITM1 (Fig. 5D; IFITM1 KO). In contrast, in the IFITM3 KO SiHa model, only a minor fraction of IFITM1 localised to the perinuclear space (Fig. 5D; white arrows), with most of the protein concentrated in patches at the plasma membrane (Fig. 5D; yellow arrows). In this cell background, only minimal diffuse cytoplasmic staining was detected.

### IFITM3 is sufficient to modulate IFITM1 localisation

The data presented above suggests that IFITM3 has the intrinsic ability to localise to perinuclear assemblies, whereas localisation of IFITM1 differs and is influenced by IFITM3 status. We therefore asked if IFITM3 was required for efficient IFITM1 perinuclear localisation and if it was sufficient to determine the subcellular distribution of IFITM1.

IFITM1:IFITM3 double knock-out E27 (Fig 5A and 6A) and E20 (Fig S1) GSCs were generated and used or transient expression of either IFITM1 or IFITM3. Using this system, we observed a mainly plasma membrane-associated localisation for exogenous IFITM1 with low levels of diffuse periplasmic staining in some cells (Fig 6A and S1, +IFITM1). On the other hand, exogenous IFITM3 localised almost exclusively to perinuclear assemblies (Fig 6A and S1, +IFITM3). Thus, exogenous IFITM3 behaves like the endogenous protein and localises to the perinuclear space regardless of IFITM1 status. When co-expressed with IFITM3, rather than being membrane associated, IFITM1 had a striking perinuclear localisation (Fig 6B; +IFITM1+IFITM3), where it was largely colocalises with IFITM3.

**Figure 6.**
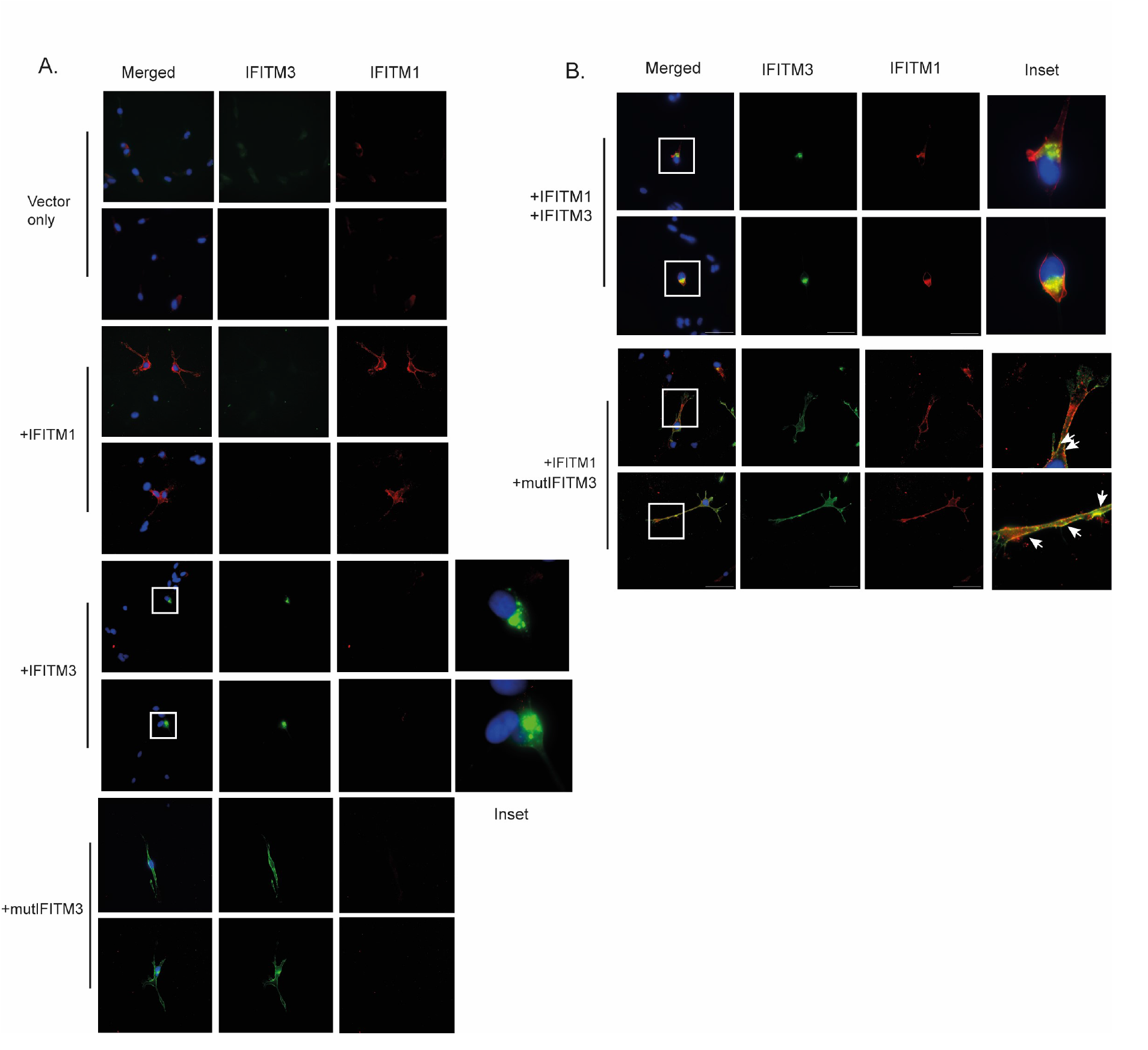
IFITM3 is sufficient to localise IFITM1 to the perinuclear space. (**A** and **B**) E27 DKO GSC cells were electroporated with empty vector or constructs expressing IFITM1, IFITM3 or Y20A-IFITM3 (mutIFITM3) as shown. The cells were treated with IFNA (500 U) for 24 h. IFITM1 was detected using Alexa 549 (red) and anti-IFITM3 was detected using Alexa 488 (green) secondary antibody. The nucleus was stained with DAPI (blue), and the scale bar is 25 μm. Arrows show areas of IFITM1:mIFITM3 co-localisation at the plasma membrane.

To confirm that the localisation of IFITM3, rather than just its presence, was required to localise IFITM1 to the perinuclear assemblies, we employed an IFITM3 mutant protein (Y20A-IFITM3). This mutant was chosen because the substitution of the Tyr residue with Ala blocks IFITM3 endocytosis, causing the protein to be retained at the plasma membrane (Fig 6A; +mutIFITM3). In the presence of mutIFITM3, IFITM1 remains at the plasma membrane and shows a degree of co-localisation with the mutIFITM3 protein (Fig 6B; +IFITM1 +mutIFITM3). Thus, we conclude that IFITM3 is required and is sufficient to promote the stable perinuclear localisation of IFITM1.

## Discussion

Investigating IFITM1 spatial organisation at both the cellular and subcellular levels within the tissue environment revealed distinct patterns in tumour endothelial cells (TECs) versus cancer stem cells in GBM tissue. Specifically, we observed that IFITM1 and IFITM3 primarily localise to the plasma membrane in glioma endothelial cells and were perinuclear in a subpopulation of tumour cells that were generally SOX2-positive (Fig 1 and 2). Using cell systems to dissect the mechanism of IFITM1 dynamics, we discovered that IFITM3 is required for efficient localisation of IFITM1 to perinuclear assemblies and that loss of IFITM3 function, or expression of a plasma membrane-associated IFITM3 mutant, leads to the accumulation of both endogenous and exogenous IFITM1 at the plasma membrane. The influence of IFITM3 on the localisation of IFITM1 highlights a regulatory mechanism that could potentially be manipulated to better understand the biochemical activity of IFITM1 in cancer whilst also providing a potential route to intervene in IFITM1 function, for example in treatment resistance and/or angiogenesis as a template for novel glioma therapeutic approaches.

How the subcellular localisation of IFITM3 links to its activity as an inhibitor of specific virus is well documented (Xie et al. 2024; Jia et al. 2012). IFITM3 is generally reported as being in the late endosomes, with recent studies showing that it impacts membrane curvature and rigidity (Guo et al. 2021; Chesarino et al. 2017) stimulating the fusion of virus bearing endosomes with lysosomes (Rahman et al. 2025). IFITM1, on the other hand, is largely described as a plasma membrane-associated protein (Smith et al. 2019; Sun et al. 2020; Weston et al. 2014). Mutations in the conserved intracellular loop of IFITM1 redistribute it from the plasma membrane to intracellular vesicles and result in a decrease in its antiviral activity towards the enveloped DNA virus HSV-1 (Smith et al. 2019). Other studies suggest vesicular IFITM1 is better able to suppress infection by virus such as JSRV (Li et al. 2015) and that the localisation of IFITM1 to multivesicular bodies is determined by a non-canonical sorting signal within its C-terminal domain. Thus, in the antiviral response, both the activity and specificity of IFITM1 may be governed by its subcellular distribution. These studies however rely heavily on over-expression of IFITM1 and the use of non-conservative mutations. As our studies show that the balance between IFITM1 and IFITM3 is a key determinant of IFITM1 subcellular distribution, we would caution against drawing definitive conclusions from overexpression studies that likely affect this finely tuned system. In the current study, we therefore largely focused on the use of tumour tissue and gene-edited KO cells to study the relationship between IFITM1 and IFITM3.

Relative to the known role of IFITM1 and -3 localisation in viral restriction, little has been published on the regulation of their subcellular distribution, and how this might inform studies on the role of IFITMs, in cancer and treatment resistance (Ogony et al. 2016; Wang et al. 2024; Yang et al. 2018; Liang, Li, and Zhu 2020). Our data suggests that IFITM3 is one of the primary determinants of IFITM1 subcellular distribution. The physiological relevance of the relationship between IFITM1 localisation and IFITM3 is supported by GBM tissue where it is notable that whilst IFITM3 is generally reported to be in the late endosomes (Jia et al. 2012; Klein et al. 2023), in glioma TECs it was at the plasma membrane where it was co-localised with IFITM1. The absence of SOX2 expression in the TECs indicates that the blood vessels were likely differentiated and mature. When we analysed established endothelial cells from the BBB, IFITM1 and -3 were not localised at the plasma membrane but were in the perinuclear space where IFITM3 localisation is expected. This suggests that signals within the local microenvironment affect the localisation of IFITM1 and -3 in glioma TECs ensuring that both proteins are retained at the plasma membrane. Alternatively, as it is emerging that TECs differ significantly from normal tissue endothelial cells, the location of IFITMs at the plasma membrane may be TEC-intrinsic (Klein et al. 2023). Consistent with roles for IFITM1 as a driver of angiogenesis through the stimulation of tube formation, it is detected at high levels in GBM tissue TECs (Fig 1), a cell type that is known for being highly angiogenic. In addition, TECs are thought to play a role in maintaining an immunosuppressive environment, which aligns with the role of IFITM1 within the ISG-RS, which is also central to the maintenance of an immunosuppressive environment (Ileiwat et al. 2022; Zeng et al. 2023).

Endocytosis of IFITM3 is regulated by phosphorylation at Y20 within an AP-2 sorting motif, ^20^YEML^23^, a region of IFITM3 that is not conserved in IFITM1 (IFITM1 has a truncated N-terminal domain relative to IFITM3). Phosphorylation at Y20 by Fyn kinase, a member of the Src family of tyrosine kinases, blocks AP-2 binding and endocytosis, leading to accumulation of IFITM3 at the plasma membrane (Chesarino et al. 2014). IFITM3-dependent antiviral functions seem to be largely restricted to its endosomal location and its ability to stimulate endosomal fusion (Spence et al. 2019). However, in B cells, IFITM3 is found on the plasma membrane where it amplifies PI3K signalling (Lee et al. 2020). A recent study provided evidence for an IFITM3 role in GBM angiogenesis (Xiong et al. 2024). However, this study focused on IFITM3 produced in GSCs and its ability to upregulate FGF secretion via activation of JAK2:STAT3 signalling. Thus, secretion of FGF into the tumour microenvironment was speculated to enhance angiogenesis in the longer term and the effects of IFITM3 in this model were not endothelial cell intrinsic. Here, we find that IFITM3 is enriched at the membrane of TECs suggesting it has a more direct role in angiogenesis. IFITM1 for example, is required for multicellular lumen formation and subsequent vessel maturation (Popson et al. 2014). In the Popson et al. (2014) study, IFITM1 is shown to regulate the expansion and maturation of nascent EC lumens through effects on the formation and/or stability of endothelial tight junctions. Therefore, it is possible to speculate that IFITM3 within the endothelial cell membrane of GBM tissue could function in cell signalling or work together with IFITM1 to regulate EC lumen development.

Whilst the role of IFITMs in cancer stem cells is not well studied, their role in ESCs (embryonic stem cells) and adult stem cells has been addressed in more depth. Thus, specific subsets of ISGs are upregulated at the RNA and protein levels in a range of stem cells (Wu et al. 2018). Of these, IFITM1 and IFITM3 appear to be key components of a stem cell ISG-signature as they are expressed at high levels in human ESCs, iPSC and tissue stem cells such as those from the hematopoietic system as well as neuronal stem cells. A role for IFITM1 and -3 in stem cells is highlighted by studies suggesting that whilst the loss of IFITMs in ESCs does not affect pluripotency, it is sufficient to overcome an innate resistance to viral infection (Wu et al. 2018). IFITM protein levels decrease during stem cell differentiation and the cells become permissive for viral replication. In parallel, whereas ISG expression in ESCs is refractive to IFNs, following differentiation, for example to hepatic-like cells, sensitivity to IFN is restored by IFITM1 attenuation. Though less is known about the role of IFITM1 and -3 in cancer stem cells (CSC), some studies provide evidence that IFITM1 is involved in CSC maintenance and regulation. For example, studies linking IFITM1 and the stem cell-associated transcription factor SOX2 are emerging. In this scenario, IFITM1 enhances the activation of the EGFR, leading to increased expression of SOX2. In NSCLC (non-small cell lung cancer), this IFITM1-mediated EGFR/SOX2 signalling axis has been shown to promote both tumour growth and metastasis (Yang et al. 2019). Both IFIMT1 and IFITM3 also appears to have a role(s) in CSC maintenance as they impact on sphere-formation, a major hallmark of stem cells (Pastrana, SilvaVargas, and Doetsch 2011). Depletion of IFITM1 reduces the ability of NSCLC cells to form spheres as well as impacting migration and invasion (Yang et al. 2019), whilst in gastric cancer cells, IFITM3 can promote cancer stemness alongside tumour growth, metastasis and chemoresistance (Chu et al. 2022).

One question raised by the results presented here is how the localisation of IFITM1 to perinuclear vesicles in tissue GSCs, rather than the plasma membrane, impacts its molecular function? Like IFITM3, IFITM1 can alter membrane fluidity and curvature, affecting processes like membrane fusion (Lin et al. 2013). However, it is not obvious how this would link to the regulation of translation (Gomez-Herranz et al. 2019; Lee et al. 2018) or activation of signalling (Wang et al. 2024; Xu, Yang, and Hu 2009). An alternative or additional possibility is that internalisation of IFITM1 is linked to endosomal signalling, a process where activated receptors and signalling molecules that have been internalised from the cell membrane, continue to function, for example, by providing a platform for the assembly of signalling complexes (Schiefermeier, Teis, and Huber 2011; Murphy et al. 2009).

In diseases like cancer, identifying how the subcellular distribution of a potential drug target varies with cell type can provide valuable information on how best to approach the development of therapeutics. For example, the location of IFITM1 on the membrane of TECs may offer an opportunity to develop IFITM1-specific antibodies aiming either to target its function in angiogenesis or to use IFITM1 to direct the delivery of anti-cancer drugs to the vasculature. On the other hand, if the primary localisation of IFITM1, for example in GSCs, suggests a role in endosomal signalling, then targeted protein degradation (e.g. PROTAC) may be a more suitable targeting approach. Thus, understanding IFITM1 dynamics can guide drug design aiming to address drug resistance mechanisms.

## Methods and Materials

### Cell culture

SiHa cells were maintained in RPMI medium (21875-034, Gibco) supplemented with 10% foetal bovine serum (FBS, Gibco) and 1% Penicillin Streptomycin (15140-122, Invitrogen).

GSCs were cultured in Dulbecco’s Modified Eagle Medium and Ham’s F12 (DMEM/F12; D8437, Merck) supplemented with 1.45 g/L D-Glucose (G8644, Merck), 120 μg/mL bovine serum albumin (BSA) fraction V (15260-037, Gibco), 100 μM β-mercaptoethanol (31350-010, Gibco), 1X MEM nonessential amino acids (MEM-NEAA; 11140-035, Gibco), 1% Penicillin Streptomycin, 0.5x B-27 (17504044, Gibco), 0.5x N-2 (15140-122, Gibco), 10 ng/mL murine EGF (315-09, Peprotech), 10 ng/mL human b-FGF (100-18b, Peprotech), and 1 μg/mL laminin-I (3446-005-01, R&D Systems).

Human cortical microvessel endothelial cells (HCMEC/D3) were cultured in Endothelial Cell Growth Medium MV2 (C-22022, PromoCell), supplemented with human recombinant epidermal growth factor (rEGF), fibroblast growth factor (rFGF), insulin-like growth factor, vascular endothelial growth factor 165, ascorbic acid and hydrocortisone. Prior to seeding, culture flasks were coated with 0.2% gelatin (G1890, Merck) and 10 µg/ml fibronectin (F1141, Merck).

All cell lines were maintained at 37 °C in a humidified incubator with 5% CO_2_ and were routinely tested for mycoplasma contamination. Cell counts were obtained using the Luna automated cell counter (Logos Biosystems) with Trypan blue exclusion.

### Immunofluorescence

Cells were washed twice with PBS, then fixed with ice-cold 100% methanol at 4 °C for 10 minutes, followed by permeabilisation in PBST (PBS + 0.1% Triton X-100) for another 10 minutes. Samples were incubated with a blocking solution (5% donkey serum in PBST) at room temperature (RT) for 30 minutes. Samples were incubated overnight at 4 °C with antibodies against IFITM1 (60074-1-Ig, Proteintech, 1:1000), IFITM3 (59212, Cell Signaling Technology, 1:1000) in blocking solution. The next day, slides were washed three times with PBST, then incubated for 1 h at room temperature with secondary antibodies - Donkey anti-Mouse IgG Alexa Fluor 594 (A21203, Invitrogen, 1:1000) and Donkey anti-Rabbit IgG Alexa Fluor 488 (A21206, Invitrogen, 1:1000) in blocking buffer. This was followed by a 5-minute incubation in 4’,6-diamidino-2-phenylindole (DAPI) (0.5 µg/ml in PBS) at room temperature after three additional PBST washes. Coverslips were mounted with Vectashield antifade mounting medium, and images were captured on a Nikon A1R+ confocal microscope. Image analysis was performed using Fiji software.

### TMA construction and slide preparation

Formalin-fixed paraffin-embedded (FFPE) glioma tissue blocks from individual patients were used to construct a tissue microarray (TMA) at the University of Edinburgh’s pathology department, as previously described (Al Shboul et al. 2024), under ethical approval provided by the Lothian NRS Bioresource (REC reference 20/ES/0061). The TMA and FFPE tissue blocks were sectioned into 3–5 μm slices using a bench-mounted microtome (ThermoFisher Scientific), and the sections were mounted onto positively charged slides (ThermoFisher Scientific) for immunohistochemistry (IHC) and immunofluorescence experiments.

### Immunohistochemistry staining

Immunohistochemistry was performed using a BOND III autostainer and the Bond Polymer Refine Detection Kit (DS9800, Leica Biosystems), following the manufacturer’s instructions. Briefly, the slides were blocked with 3–4% (v/v) hydrogen peroxide for 5 minutes. Antigen retrieval was performed using Bond Epitope Retrieval Solution 1 (Citrate, pH 6.0) (AR9961, Leica Biosystems) for 20 minutes at 100°C. The slides were then incubated at room temperature for 20 minutes with the antibodies against IFITM1 (60074-1-Ig, Proteintech, 1:1000), IFITM3 (59212, Cell Signaling Technology, 1:200), and SOX2 (ab97959, Abcam, 1:200). Detection was achieved using 3,3′diaminobenzidine (DAB) staining and haematoxylin counterstaining (Leica Biosystems). After staining, the slides were dehydrated, cleared with xylene, and coverslips were applied. The TMA slides were scanned at 40x magnification using a NanoZoomer XR Digital Pathology slide scanner (Hamamatsu), and images were analysed using QuPath (version 0.3.1).

### Tissue immunofluorescence

Slides prepared from FFPE tissue blocks were dewaxed twice in xylene for 3 minutes and rehydrated through a series of ethanol washes (twice in 100%, 90%, and 70% ethanol for 3 minutes each). The slides were then stored in distilled water for 15 minutes. Freshly prepared antigen retrieval buffer (0.1 M sodium citrate buffer, pH 6.0, with 0.5% Tween 20) was heated for 10 minutes in a pressure cooker at an internal temperature of 90–125°C. Slides were immersed in the buffer and heated for an additional 5 minutes. After antigen retrieval, the buffer was cooled to room temperature, and the slides were washed with PBS. Next, the slides were incubated in the blocking buffer (5% BSA with 0.2% Triton X-100 in Tris-buffered saline (TBS)) for 1 h at RT. They were then incubated overnight at 4°C with primary antibodies against IFITM1 (60074-1-Ig, Proteintech, 1:1000), IFITM3 (59212, Cell Signaling Technology, 1:1000), CD31 (ab28364, Abcam, 1:100), and SOX2 (ab97959, Abcam, 1:200). The following day, after three washes with TBST, the slides were incubated for 1 h at RT with Donkey anti-Mouse IgG Alexa Fluor 594 (A21203, Invitrogen, 1:1000) and Donkey anti-Rabbit IgG Alexa Fluor 488 secondary antibody (A21206, Invitrogen, 1:1000) in blocking buffer. After three additional washes with TBST, the slides were stained with DAPI (0.5 µg/ml in PBS) for 5 minutes at RT, mounted with Vectashield antifade mounting medium, and imaged using a Nikon A1R+ confocal microscope. The acquired images were analysed using Fiji software.

### CRISPR/Cas9 Gene Editing

IFITM1 and IFITM3 KO SiHa cells were generated either as previously described (Gomez-Herranz et al. 2019) or using ribonucleoprotein (RNP)-based method with lipofectamine CRISPRMAX Cas9 transfection reagent (CMAX00001, Invitrogen). Guide RNAs (gRNAs) targeting IFITM1 or IFITM3 exons were ordered as crRNAs from IDT. RNP complexes were prepared by combining 100 pmol of each crRNA and tracrRNA in IDT resuspension buffer to a final volume of 100 µL, followed by denaturation at 95 °C for 5 minutes and gradual cooling for annealing. Subsequently, recombinant Cas9 protein was diluted to a final concentration of 1 µM in Opti-MEM (Gibco). For each well of a 6well plate, 38.75 µL of the annealed gRNA:tracrRNA duplex was combined with 38.75 µL Cas9 and 35 µL Opti-MEM, followed by the addition of 12.5 µL Cas9 Plus Reagent, yielding a total volume of 125 µL. This mixture was incubated at RT for 5 minutes. In a separate tube, 7.5 µL of CRISPRMAX reagent was diluted in 117.5 µL Opti-MEM and immediately combined with the RNP complex. The final transfection mix was incubated for 15-30 minutes at RT. Meanwhile, 200,000 SiHa cells/ml were plated into each well of a 6-well plate in 2.25 ml of antibiotic-free complete RPMI medium. Subsequently, 250 µL of the prepared transfection complex was added to the cells and incubated at 37 °C with 5% CO_2_ for 48 hours. Following recovery, cells were passaged once prior to single-cell sorting by flow cytometry.

IFITM1 and IFITM3 KO GSCs were generated as previously described (Dewari et al. 2018). Briefly, 100 pmol of each crRNA and tracrRNA were mixed, denatured at 95°C, and gradually cooled in a thermocycler to allow annealing. 10 µg recombinant Cas9 protein was added to annealed cr/tracrRNAs and incubated at RT for 10 minutes to form the RNP complex. This RNP complex was then electroporated into 2×10^5^ cells using the Nucleofector Kit SG (197175, Lonza) with the DN-100 program. Cells were allowed to recover for 48-72 hours prior to single-cell sorting by flow cytometry.

Further, the cells were trypsinised and resuspended in 5 ml of 0.2% FBS in 1x sterile PBS and single cells were sorted in each well of 96-well plates using flow cytometry. Individual colonies were further propagated and screened for IFITM1 and IFITM3 expression with interferon stimulation using western blotting, PCR, followed by Sanger sequencing (SourceBioscience) to obtain specific gene edits.

### Transient transfection of plasmid DNA

Cells were trypsinised and washed twice with Opti-MEM (Gibco, 31985062). An appropriate number of cells was resuspended in Opti-MEM, 5 µg of plasmid DNA – either pcDNA3.1-IFITM1, pcDNA3.1-IFITM3 Y20A or pSF-CMV-FLuc-CMV-Zep-BgH-sbf1-IFITM3 was added to a final volume of 100 µL. The mixtures were transferred to NEPA nucleofection cuvettes, and nucleofection was performed using a NEPA nucleofector with the following parameters: poring pulse at 125 V and transfer pulse at 20 V. Immediately after nucleofection, 400 µL of pre-warmed medium was added, and cells were plated onto laminin-coated 12-well plates containing coverslips for subsequent immunofluorescence microscopy.

### Flp-In cell lines

To generate IFITM1-expressing Flp-In stable cell lines, the Flp-In complete system (K601001; ThermoFisher Scientific) was used according to the manufacturer’s instructions. Briefly, Flp-In host cell lines were first established in IFITM1 KO cells by transfecting the pFRT/lacZeo plasmid, which carries the FRT recombination site, using Attractene transfection reagent (Qiagen) as per the manufacturer’s protocol. Zeocin-resistant colonies were selected and maintained in medium containing 50 µg/mL Zeocin (Thermofisher Scientific) until co-transfection with pOG44 and PCDNA5/FRT plasmids. Successful integration of the FRT site was confirmed by PCR amplification of the LacZ gene.

To generate IFITM1-expressing stable lines, Flp-In host cells were co-transfected with the pOG44 plasmid (encoding Flp recombinase) and the pcDNA5/FRT vector containing WT IFITM1. A similar procedure was performed with the pFRT/lacZeo control plasmids. Following transfection, cells were selected in hygromycin B-containing medium (Millipore), and resistant colonies were expanded to establish stable cell lines.

## Acknowledgements

We are grateful to core services staff at the Institute of Genetics and Cancer for their expertise and support. We acknowledge Matthew Pearson and James Iremonger from the Advanced Imaging Resource at the Institute of Genetics and Cancer, University of Edinburgh, for their imaging support. YSB and EN were funded by studentship awards to KLB by BBSRC/IBioIC DTP and Medical Research Scotland, respectively. AS was funded by the EU Horizon 2020 Research and Innovation program under agreement number 101017453 (KATY).

## References

Ahir, B. K., H. H. Engelhard, and S. S. Lakka. 2020. ‘Tumor Development and Angiogenesis in Adult Brain Tumor: Glioblastoma’, Mol Neurobiol, 57: 2461–78.

Al Shboul, S., S. Boyle, A. Singh, T. Saleh, M. Alrjoub, O. Abu Al Karsaneh, A. Mryyian, R. Dawoud, S. Gul, S. Abu Baker, K. Ball, T. Hupp, and P. M. Brennan. 2024. ‘FISH analysis reveals CDKN2A and IFNA14 co-deletion is heterogeneous and is a prominent feature of glioblastoma’, Brain Tumor Pathol, 41: 4–17.

Benci, J. L., L. R. Johnson, R. Choa, Y. Xu, J. Qiu, Z. Zhou, B. Xu, D. Ye, K. L. Nathanson, C. H. June, E. J. Wherry, N. R. Zhang, H. Ishwaran, M. D. Hellmann, J. D. Wolchok, T. Kambayashi, and A. J. Minn. 2019. ‘Opposing Functions of Interferon Coordinate Adaptive and Innate Immune Responses to Cancer Immune Checkpoint Blockade’, Cell, 178: 933–48 e14.

Chesarino, N. M., A. A. Compton, T. M. McMichael, A. D. Kenney, L. Zhang, V. Soewarna, M. Davis, O. Schwartz, and J. S. Yount. 2017. ‘IFITM3 requires an amphipathic helix for antiviral activity’, EMBO Rep, 18: 1740–51.

Chesarino, N. M., T. M. McMichael, J. C. Hach, and J. S. Yount. 2014. ‘Phosphorylation of the antiviral protein interferon-inducible transmembrane protein 3 (IFITM3) dually regulates its endocytosis and ubiquitination’, J Biol Chem, 289: 11986–92.

Chu, P. Y., W. C. Huang, S. L. Tung, C. Y. Tsai, C. J. Chen, Y. C. Liu, C. W. Lee, Y. H. Lin, H. Y. Lin, C. Y. Chen, C. T. Yeh, K. H. Lin, and H. C. Chi. 2022. ‘IFITM3 promotes malignant progression, cancer stemness and chemoresistance of gastric cancer by targeting MET/AKT/FOXO3/cMYC axis’, Cell Biosci, 12: 124.

Dewari, P. S., B. Southgate, K. McCarten, G. Monogarov, E. O’Duibhir, N. Quinn, A. Tyrer, M. C. Leitner, C. Plumb, M. Kalantzaki, C. Blin, R. Finch, R. B. Bressan, G. Morrison, A. M. Jacobi, M. A. Behlke, A. von Kriegsheim, S. Tomlinson, J. Krijgsveld, and S. M. Pollard. 2018. ‘An efficient and scalable pipeline for epitope tagging in mammalian stem cells using Cas9 ribonucleoprotein’, Elife, 7.

Eckerdt, F., and L. C. Platanias. 2023. ‘Emerging Role of Glioma Stem Cells in Mechanisms of Therapy Resistance’, Cancers (Basel), 15.

Friedlova, N., F. Zavadil Kokas, T. R. Hupp, B. Vojtesek, and M. Nekulova. 2022. ‘IFITM protein regulation and functions: Far beyond the fight against viruses’, Front Immunol, 13: 1042368.

Gomez-Herranz, M., M. Nekulova, J. Faktor, L. Hernychova, S. Kote, E. H. Sinclair, R. Nenutil, B. Vojtesek, K. L. Ball, and T. R. Hupp. 2019. ‘The effects of IFITM1 and IFITM3 gene deletion on IFNgamma stimulated protein synthesis’, Cell Signal, 60: 39–56.

Gomez-Herranz, M., J. Taylor, and R. D. Sloan. 2023. ‘IFITM proteins: Understanding their diverse roles in viral infection, cancer, and immunity’, J Biol Chem, 299: 102741.

Guo, X., J. Steinkuhler, M. Marin, X. Li, W. Lu, R. Dimova, and G. B. Melikyan. 2021. ‘InterferonInduced Transmembrane Protein 3 Blocks Fusion of Diverse Enveloped Viruses by Altering Mechanical Properties of Cell Membranes’, ACS Nano, 15: 8155–70.

Ileiwat, Z. E., T. A. Tabish, D. A. Zinovkin, J. Yuzugulen, N. Arghiani, and M. Z. I. Pranjol. 2022. ‘The mechanistic immunosuppressive role of the tumour vasculature and potential nanoparticlemediated therapeutic strategies’, Front Immunol, 13: 976677.

Jia, R., Q. Pan, S. Ding, L. Rong, S. L. Liu, Y. Geng, W. Qiao, and C. Liang. 2012. ‘The N-terminal region of IFITM3 modulates its antiviral activity by regulating IFITM3 cellular localization’, J Virol, 86: 13697–707.

Klein, S., G. Golani, F. Lolicato, C. Lahr, D. Beyer, A. Herrmann, M. Wachsmuth-Melm, N. Reddmann, R. Brecht, M. Hosseinzadeh, A. Kolovou, J. Makroczyova, S. Peterl, M. Schorb, Y. Schwab, B. Brugger, W. Nickel, U. S. Schwarz, and P. Chlanda. 2023. ‘IFITM3 blocks influenza virus entry by sorting lipids and stabilizing hemifusion’, Cell Host Microbe, 31: 616–33 e20.

Lee, J., M. E. Robinson, N. Ma, D. Artadji, M. A. Ahmed, G. Xiao, T. Sadras, G. Deb, J. Winchester, K. N. Cosgun, H. Geng, L. N. Chan, K. Kume, T. P. Miettinen, Y. Zhang, M. A. Nix, L. Klemm, C. W. Chen, J. Chen, V. Khairnar, A. P. Wiita, A. Thomas-Tikhonenko, M. Farzan, J. U. Jung, D. M. Weinstock, S. R. Manalis, M. S. Diamond, N. Vaidehi, and M. Muschen. 2020. ‘IFITM3 functions as a PIP3 scaffold to amplify PI3K signalling in B cells’, Nature, 588: 491–97.

Lee, W. J., R. M. Fu, C. Liang, and R. D. Sloan. 2018. ‘IFITM proteins inhibit HIV-1 protein synthesis’, Sci Rep, 8: 14551.

Li, K., R. Jia, M. Li, Y. M. Zheng, C. Miao, Y. Yao, H. L. Ji, Y. Geng, W. Qiao, L. M. Albritton, C. Liang, and S. L. Liu. 2015. ‘A sorting signal suppresses IFITM1 restriction of viral entry’, J Biol Chem, 290: 4248–59.

Liang, R., X. Li, and X. Zhu. 2020. ‘Deciphering the Roles of IFITM1 in Tumors’, Mol Diagn Ther, 24: 433–41.

Lin, T. Y., C. R. Chin, A. R. Everitt, S. Clare, J. M. Perreira, G. Savidis, A. M. Aker, S. P. John, D. Sarlah, E. M. Carreira, S. J. Elledge, P. Kellam, and A. L. Brass. 2013. ‘Amphotericin B increases influenza A virus infection by preventing IFITM3-mediated restriction’, Cell Rep, 5: 895–908.

Liu, Z. L., H. H. Chen, L. L. Zheng, L. P. Sun, and L. Shi. 2023. ‘Angiogenic signaling pathways and antiangiogenic therapy for cancer’, Signal Transduct Target Ther, 8: 198.

Mosteiro, A., L. Pedrosa, A. Ferres, D. Diao, A. Sierra, and J. J. Gonzalez. 2022. ‘The Vascular Microenvironment in Glioblastoma: A Comprehensive Review’, Biomedicines, 10.

Murphy, J. E., B. E. Padilla, B. Hasdemir, G. S. Cottrell, and N. W. Bunnett. 2009. ‘Endosomes: a legitimate platform for the signaling train’, Proc Natl Acad Sci U S A, 106: 17615–22.

Ogony, J., H. J. Choi, A. Lui, M. Cristofanilli, and J. Lewis-Wambi. 2016. ‘Interferon-induced transmembrane protein 1 (IFITM1) overexpression enhances the aggressive phenotype of SUM149 inflammatory breast cancer cells in a signal transducer and activator of transcription 2 (STAT2)-dependent manner’, Breast Cancer Res, 18: 25.

Pastrana, E., V. Silva-Vargas, and F. Doetsch. 2011. ‘Eyes wide open: a critical review of sphereformation as an assay for stem cells’, Cell Stem Cell, 8: 486–98.

Popson, S. A., M. E. Ziegler, X. Chen, M. T. Holderfield, C. I. Shaaban, A. H. Fong, K. M. WelchReardon, J. Papkoff, and C. C. Hughes. 2014. ‘Interferon-induced transmembrane protein 1 regulates endothelial lumen formation during angiogenesis’, Arterioscler Thromb Vasc Biol, 34: 1011–9.

Rahman, K., I. Wilt, A. A. Jolley, B. Chowdhury, S. A. K. Datta, and A. A. Compton. 2025. ‘SNARE mimicry by the CD225 domain of IFITM3 enables regulation of homotypic late endosome fusion’, EMBO J, 44: 534–62.

Rahman, M. A., and M. M. Ali. 2024. ‘Recent Treatment Strategies and Molecular Pathways in Resistance Mechanisms of Antiangiogenic Therapies in Glioblastoma’, Cancers (Basel), 16.

Rajapaksa, U. S., C. Jin, and T. Dong. 2020. ‘Malignancy and IFITM3: Friend or Foe?’, Front Oncol, 10: 593245.

Schiefermeier, N., D. Teis, and L. A. Huber. 2011. ‘Endosomal signaling and cell migration’, Curr Opin Cell Biol, 23: 615–20.

Shi, T., J. Zhu, X. Zhang, and X. Mao. 2023. ‘The Role of Hypoxia and Cancer Stem Cells in Development of Glioblastoma’, Cancers (Basel), 15.

Smith, S. E., D. C. Busse, S. Binter, S. Weston, C. Diaz Soria, B. M. Laksono, S. Clare, S. Van Nieuwkoop, B. G. Van den Hoogen, M. Clement, M. Marsden, I. R. Humphreys, M. Marsh, R. L. de Swart, R. S. Wash, J. S. Tregoning, and P. Kellam. 2019. ‘Interferon-Induced Transmembrane Protein 1 Restricts Replication of Viruses That Enter Cells via the Plasma Membrane’, J Virol, 93.

Spence, J. S., R. He, H. H. Hoffmann, T. Das, E. Thinon, C. M. Rice, T. Peng, K. Chandran, and H. C. Hang. 2019. ‘IFITM3 directly engages and shuttles incoming virus particles to lysosomes’, Nat Chem Biol, 15: 259–68.

Sun, F., Z. Xia, Y. Han, M. Gao, L. Wang, Y. Wu, J. M. Sabatier, L. Miao, and Z. Cao. 2020. ‘Topology, Antiviral Functional Residues and Mechanism of IFITM1’, Viruses, 12.

Tang, X., C. Zuo, P. Fang, G. Liu, Y. Qiu, Y. Huang, and R. Tang. 2021. ‘Targeting Glioblastoma Stem Cells: A Review on Biomarkers, Signal Pathways and Targeted Therapy’, Front Oncol, 11: 701291.

Wang, X., H. Qian, L. Yang, S. Yan, H. Wang, X. Li, and D. Yang. 2024. ‘The role and mechanism of IFITM1 in developing acquired cisplatin resistance in small cell lung cancer’, Heliyon, 10: e30806.

Weichselbaum, R. R., H. Ishwaran, T. Yoon, D. S. Nuyten, S. W. Baker, N. Khodarev, A. W. Su, A. Y. Shaikh, P. Roach, B. Kreike, B. Roizman, J. Bergh, Y. Pawitan, M. J. van de Vijver, and A. J. Minn. 2008. ‘An interferon-related gene signature for DNA damage resistance is a predictive marker for chemotherapy and radiation for breast cancer’, Proc Natl Acad Sci U S A, 105: 18490–5.

Weston, S., S. Czieso, I. J. White, S. E. Smith, P. Kellam, and M. Marsh. 2014. ‘A membrane topology model for human interferon inducible transmembrane protein 1’, PLoS One, 9: e104341.

Wu, X., V. L. Dao Thi, Y. Huang, E. Billerbeck, D. Saha, H. H. Hoffmann, Y. Wang, L. A. V. Silva, S. Sarbanes, T. Sun, L. Andrus, Y. Yu, C. Quirk, M. Li, M. R. MacDonald, W. M. Schneider, X. An, B. R. Rosenberg, and C. M. Rice. 2018. ‘Intrinsic Immunity Shapes Viral Resistance of Stem Cells’, Cell, 172: 423–38 e25.

Xie, Q., L. Wang, X. Liao, B. Huang, C. Luo, G. Liao, L. Yuan, X. Liu, H. Luo, and Y. Shu. 2024. ‘Research Progress into the Biological Functions of IFITM3’, Viruses, 16.

Xiong, Z., X. Xu, Y. Zhang, C. Ma, C. Hou, Z. You, L. Shu, Y. Ke, and Y. Liu. 2024. ‘IFITM3 promotes glioblastoma stem cell-mediated angiogenesis via regulating JAK/STAT3/bFGF signaling pathway’, Cell Death Dis, 15: 45.

Xu, Y., G. Yang, and G. Hu. 2009. ‘Binding of IFITM1 enhances the inhibiting effect of caveolin-1 on ERK activation’, Acta Biochim Biophys Sin (Shanghai), 41: 488–94.

Yang, J., L. Li, Y. Xi, R. Sun, H. Wang, Y. Ren, L. Zhao, X. Wang, and X. Li. 2018. ‘Combination of IFITM1 knockdown and radiotherapy inhibits the growth of oral cancer’, Cancer Sci, 109: 3115–28.

Yang, Y. G., Y. W. Koh, I. N. Sari, N. Jun, S. Lee, L. T. H. Phi, K. S. Kim, Y. T. Wijaya, S. H. Lee, M. J. Baek, D. Jeong, and H. Y. Kwon. 2019. ‘Interferon-induced transmembrane protein 1-mediated EGFR/SOX2 signaling axis is essential for progression of non-small cell lung cancer’, Int J Cancer, 144: 2020–32.

Yu, F., S. S. Ng, B. K. Chow, J. Sze, G. Lu, W. S. Poon, H. F. Kung, and M. C. Lin. 2011. ‘Knockdown of interferon-induced transmembrane protein 1 (IFITM1) inhibits proliferation, migration, and invasion of glioma cells’, J Neurooncol, 103: 187–95.

Zeng, Q., M. Mousa, A. S. Nadukkandy, L. Franssens, H. Alnaqbi, F. Y. Alshamsi, H. A. Safar, and P. Carmeliet. 2023. ‘Understanding tumour endothelial cell heterogeneity and function from single-cell omics’, Nat Rev Cancer, 23: 544–64.

