## Supplementary figure for "Spatial dynamics of IFITM1: a core component of the interferon-stimulated gene-resistance signature in glioma"

Supplementary Data:

**Figure S1:** Exogenous IFITM1 localises primarily to the plasma membrane in the absence of IFITM3. (**A**) WT and DKO E20 GSCs were treated with IFNA (500 U) or carrier alone for 24 h prior to lysis. Immunoblots were developed using anti-IFITM1 or anti-IFITM3 antibodies and the IFITM1 membrane was subsequently overlayed with anti-β-actin antibody. (**B**) Immunofluorescence of WT E20 GSCs (WT-GSC), IFITM1 and IFITM3 double KOs (DKO) and the DKO expressing exogenous IFITM1 (DKOGSC+IFITM1) or IFITM3 (DKO-GSC+IFITM3) after incubation with IFNA (500 U) for 24 h. IFITM1 was detected using Alexa 549 (red) and IFITM3 was detected using Alexa 488 (green) secondary antibody. The nucleus was stained with DAPI (blue), and the scale bar is 25 µm. Images shown are representative of two independent experiments.

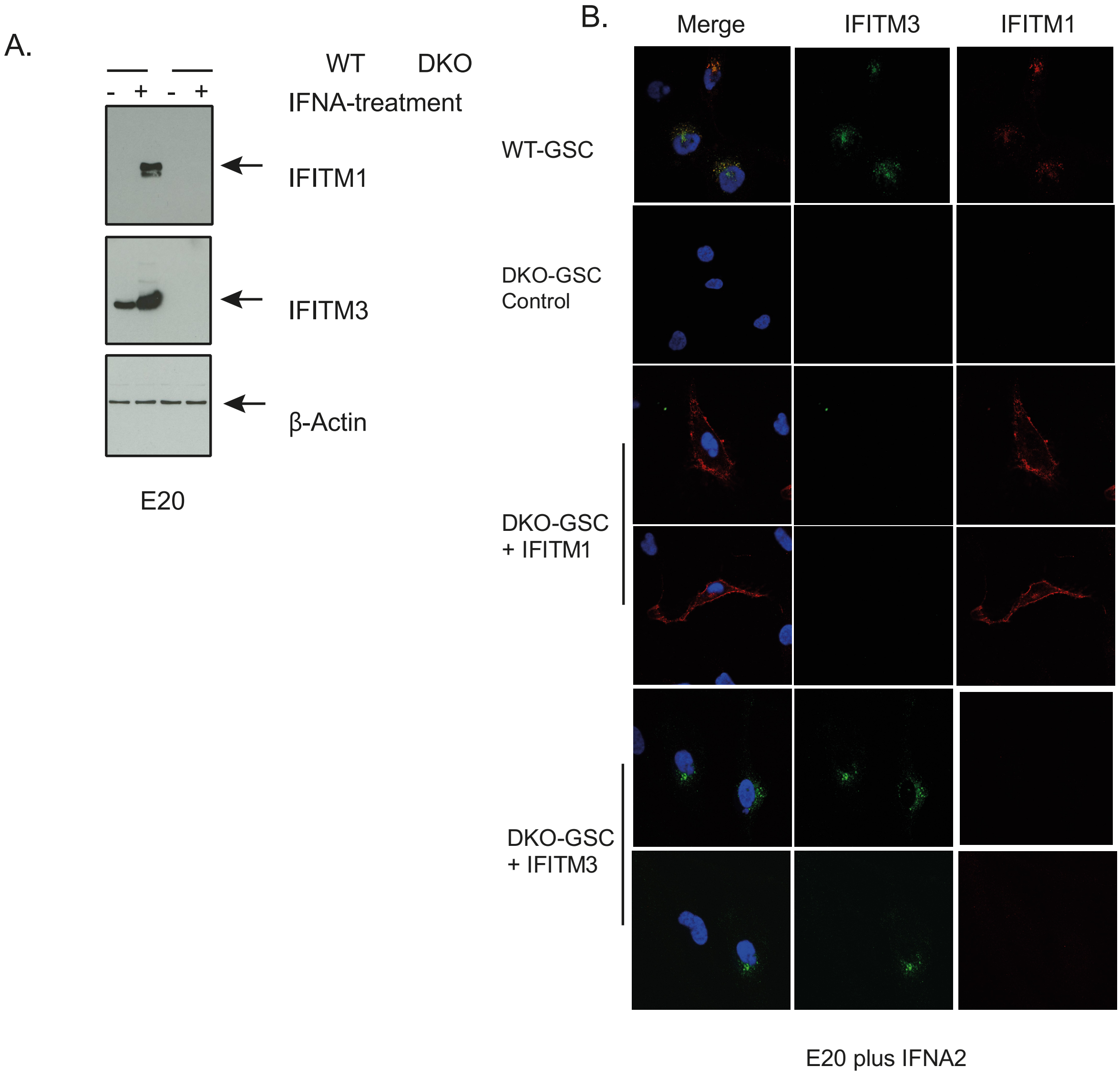
